## Supplemental Figure 1 for "Geography is a stronger predictor of diversification of monogenean parasites (Platyhelminthes) than host relatedness in characid fishes of Middle America"

**S1 Fig. Cophylogenetic plot showing associations between characid fish hosts and their monogenean parasites.** Phylogenetic trees are ML trees in Figs 1B and 2A. Line colors correspond to areas of endemism.

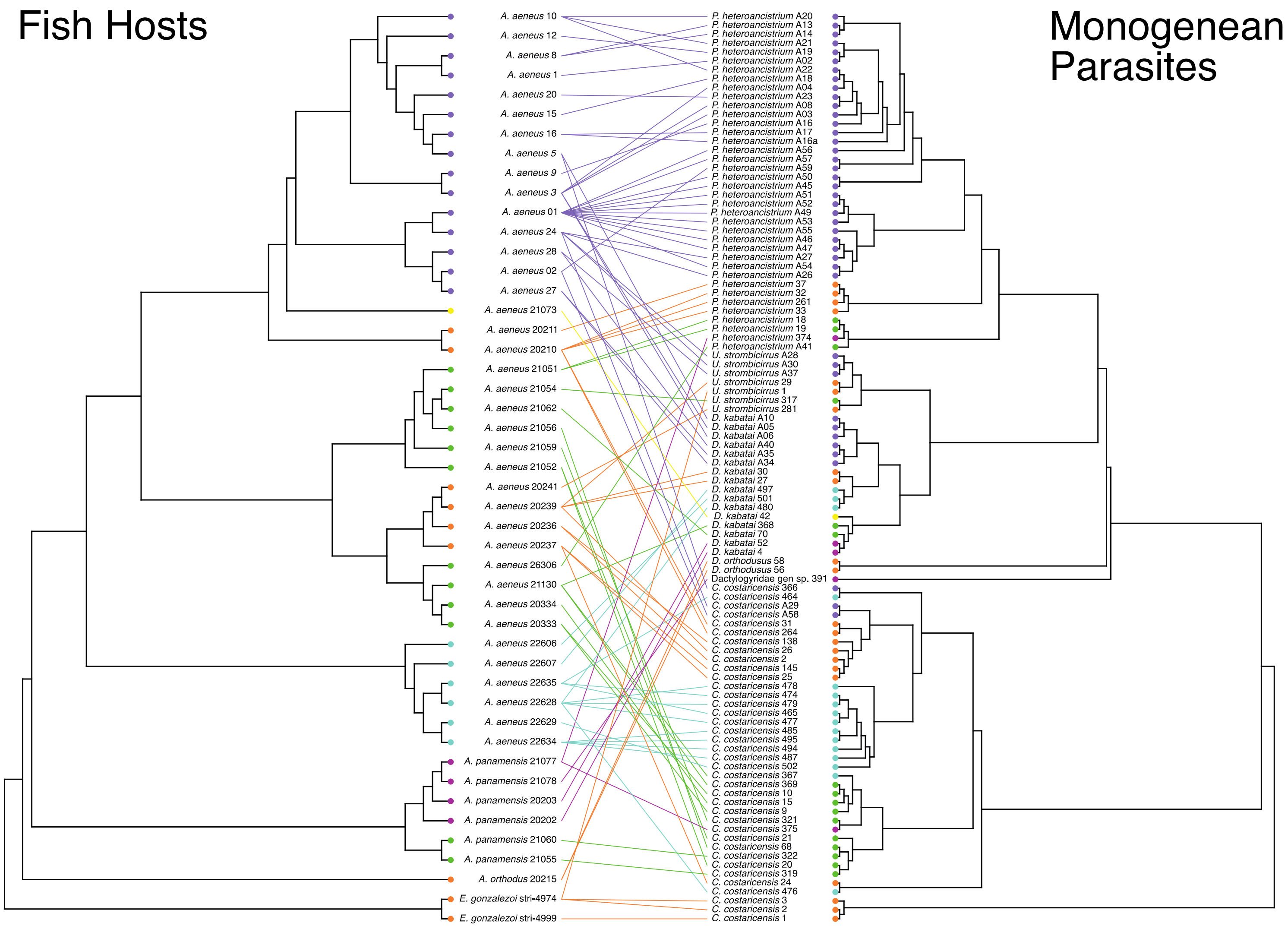
